# Prenatal exposure to PFAS and multimodal structural brain development across childhood

**DOI:** 10.64898/2026.08.13.744565

**Authors:** Sarah Rocha, Jessica P. Uy, Chase Antonacci, Jessica L. Buthmann, Ai Peng Tan, Yap Seng Chong, Marielle V. Fortier, Johan Eriksson, Ian H. Gotlib

**Affiliations:** Stanford University, Department of Psychology, Building 420, 450 Jane Stanford Way, Stanford, CA 94305, USA; Institute for Human Development and Potential, Agency for Science, Technology and Research (A*STAR), Singapore; Department of Diagnostic Imaging, National University Health System, Singapore; Yong Loo Lin School of Medicine, National University of Singapore, Singapore; KK Women and Children’s Hospital, Singapore; Duke-National University of Singapore, Singapore; National University of Singapore, National University Health System, Singapore; University of Helsinki, Helsinki, Finland; Folkhälsan Research Center, Helsinki, Finland

**Keywords:** per- and polyfluoroalkyl substances, magnetic resonance imaging, white matter microstructure, neurodevelopment, longitudinal

## Abstract

**Background:** Per- and polyfluoroalkyl substances (PFAS) are ubiquitous environmental pollutants that are posited to be neurotoxic to the developing brain; however, the impact of prenatal exposure to PFAS — particularly to newer, short-chain PFAS — on brain development across childhood is unclear.

**Methods:** Concentrations of 9 PFAS were quantified in cord blood plasma of 459 infants who later participated in structural and diffusion magnetic resonance imaging (MRI) at ages 4.5, 6, 7.5, and 10.5 years, providing estimates of regional cortical thickness and surface area, subcortical volumes, and fractional anisotropy (FA) of key white matter tracts. Longitudinal mixed effect models estimated associations of PFAS with age 4.5 brain metrics and their developmental trajectories across childhood.

**Results:** Higher concentrations of long-chain PFAS (PFNA, PFHxS, PFDA) in cord blood were associated with lower surface area of the right paracentral lobule at age 4.5. PFHpA was associated with faster surface area growth in the left rostral anterior cingulate and slower growth in the right caudal middle frontal gyrus from 4.5 to 10.5 years. The short-chain compound PFBS was linked with greater FA in 17 of 27 white matter tracts at 4.5 years; those associations attenuated with age. Finally, PFOA was associated with lower FA in 6 tracts at 4.5 years.

**Conclusions:** Prenatal exposure to PFAS was associated with altered development of frontal and paracentral regions and of white matter microstructure. These findings highlight the need for further research examining the long-term effects of prenatal exposure to PFAS on children’s neurodevelopment.

## Introduction

Per- and polyfluoroalkyl substances (PFAS) are highly stable, synthetically fluorinated chemicals that are among the most widespread environmental contaminants globally (Cousins et al. 2022). PFAS have been used by manufacturers since the early 1950s because their chemical structure confers resistance to degradation and also provides water- and oil-repelling properties (Gaines 2023). Today, PFAS are used widely in the production of a wide range of industrial and consumer products, such as nonstick cookware, water-proof clothing, food packaging, building materials, and firefighting foams (Glüge et al. 2020). While certain long-chain “legacy” PFAS such as perfluorooctanoic acid (PFOA) and perfluorooctanesulfonic acid (PFOS) have been phased out of production following evidence of harmful effects on human reproductive health (Chambers, Hopkins, and Richards 2021), endocrine function (Coperchini et al. 2021), and risk for cancer (Steenland and Winquist 2021), they continue to accumulate in the environment due to their long half-lives (Cousins et al. 2022). Importantly, they are also being replaced by newer, short-chain PFAS for which effects on human health are not well understood (Kannan et al. 2026).

Humans are exposed to PFAS in multiple ways, including ingestion of PFAS-contaminated water and food, dermal contact with consumer products containing PFAS, and inhalation of airborne emissions of PFAS (Sunderland et al. 2019). These many routes of exposure have resulted in near ubiquitous detection of PFAS in humans (Botelho et al. 2025), including in such sensitive groups as pregnant individuals and infants (Kirk et al. 2022). There is increasing evidence that exposure to PFAS during gestation has adverse neurodevelopmental consequences. Most notably, prenatal exposure to PFAS has been associated with developmental delays in language and motor skills (Luo et al. 2022), attention problems and attention-deficit hyperactivity disorder (ADHD; Kim et al. 2023), childhood behavioral problems (Tillaut et al. 2023), and a higher incidence of autism spectrum disorder (Oh et al. 2021). The prenatal period is a window of rapid fetal brain development and heightened sensitivity to environmental contaminants (Gómez-Roig et al. 2021). PFAS can be transferred from the mother to the developing fetus through the placenta (Gao et al. 2019) and are also able to cross the blood-brain barrier (Liu et al. 2022) and accumulate in cerebrospinal fluid and brain tissue. Researchers have posited that PFAS may adversely affect brain development by impairing calcium homeostasis in neurons and disrupting the secretion of such neurotransmitters as dopamine and glutamate (Brown-Leung and Cannon 2022). PFAS may also induce neurotoxicity indirectly by dysregulating endocrine functions that influence neural development and by increasing inflammatory signaling (Lagostena et al. 2025).

Despite the ubiquitous presence of PFAS in the environment and their links with children’s neurodevelopmental deficits, few studies have examined the associations of prenatal exposure to PFAS with children’s brain development. These investigations, which are typically cross-sectional and small in scale (i.e., samples of 49–155 children), have yielded mixed findings. For example, in 49 Taiwanese adolescents, greater prenatal exposure to specific long-chain PFAS (i.e., PFOS, perfluorononanoic acid [PFNA] and perfluorododecanoic acid [PFDoA]) was associated with smaller volume of the frontal lobe and cerebellum (Shen et al. 2021); further, PFDoA was linked with reduced fiber integrity of two white matter tracts (right superior and inferior cerebellar peduncles; Wang et al. 2025). Two other studies have also documented associations of prenatal exposure to PFAS with altered white matter microstructure, although in opposite directions. Whereas England-Mason et al. (2025) found that prenatal exposure to PFOS and perfluorohexanesulfonic acid (PFHxS) was associated with reduced fiber integrity of the corpus callosum in a Canadian cohort of 84 children ages 2-6 years, Barron et al. (2025) documented associations between PFNA and PFOA and a multimodal neuroimaging component score dominated by *greater* integrity of the corpus callosum in a Finnish cohort of 51 5-year-old children. Finally, Haraburda et al. (2026) found no associations of prenatal exposure to PFAS with metrics of gray or white matter structure in 12-year-old children.

Although these studies provide preliminary evidence that prenatal exposure to PFAS may influence children’s structural brain development, particularly the fiber integrity of white matter tracts, there are two issues that warrant consideration. First, these studies did not examine the effects of newer, short-chain PFAS on children’s brain structure. To comply with international regulations of certain long-chain PFAS (e.g., PFOA, PFOS, PFNA), manufacturers are increasingly using short-chain “replacement” PFAS such as perfluorobutanesulfonic acid (PFBS) and perfluorobutanoic acid (PFBA) in consumer and industrial products. Although these short-chain PFAS were thought to have lower toxicity due to their shorter biological half-lives, there is now evidence that they, too, may have neurotoxic effects (Wasel et al. 2023), which is particularly worrisome given that their higher water solubility and motility allow for wider dispersal (Kannan et al. 2026). Importantly, short-chain PFAS are becoming increasingly prevalent across global environments (Kannan et al. 2026), with particularly high levels of PFBS in Singapore (Chen et al. 2017), from where we are analyzing data for the present study. Second, given the cross-sectional designs of previous studies, we know little about the effects of prenatal exposure to PFAS on the *development* of children’s brain structure. Given that researchers have documented dynamic changes in gray and white matter structure from early childhood to adolescence (Lebel, Treit, and Beaulieu 2019; Mills et al. 2016), it is critical that we elucidate the effects of prenatal PFAS exposure on the trajectory of brain structure across this period.

The present study was designed to characterize the associations of prenatal exposure to both long- and short-chain PFAS on brain development longitudinally across childhood. Specifically, we leveraged multimodal neuroimaging data from *N*=459 children from the longitudinal birth cohort study, *Growing up in Singapore towards healthy Outcomes* (GUSTO). The present study is the largest, multimodal investigation of prenatal PFAS and structural brain development to date, and the only study to examine longitudinal trajectories of multimodal brain development across childhood. Concentrations of 9 PFAS were estimated from infant umbilical cord blood, and children subsequently completed up to four structural and diffusion magnetic resonance imaging (MRI) scans across the first decade of life (at 4.5, 6, 7.5, and 10.5 years of age). These scans were used to compute estimates of cortical thickness and surface area, subcortical volumes, and fractional anisotropy (FA) of major white matter tracts at each of these four timepoints. Based on previous findings (Shen et al. 2021), we hypothesized, first, that higher prenatal exposure to long-chain PFAS will be associated with lower cortical thickness and surface area of frontal brain regions. Second, we hypothesized this prenatal exposure will also be associated with smaller volumes of subcortical structures in which higher levels of PFAS are thought to accumulate, such as the thalamus, hippocampus, basal ganglia, and brain stem (Di Nisio et al. 2022). Third, we hypothesized that prenatal exposure to long-chain PFAS will be associated with alterations in white matter microstructure; given conflicting findings in previous research (Barron et al. 2025; England-Mason et al. 2025), we did not make this hypothesis directional. Finally, given the lack of prior studies, we also did not generate directional hypotheses about the associations of prenatal exposure to short-chain PFAS with trajectories of brain development.

## Methods

### Participants

Participants were drawn from the ongoing, longitudinal birth-cohort study, Growing Up in Singapore Towards healthy Outcomes (GUSTO). The cohort has been described in detail previously (Soh et al. 2014). Briefly, pregnant mothers were recruited from public maternity hospitals in Singapore from 2009 to 2010. Mothers were eligible to participate if they were planning to deliver at the maternity hospital and continue to reside in Singapore for at least five years, and if they agreed to donate cord tissue following delivery; ineligibility criteria included receiving chemotherapy, use of psychotropic drugs and having type I diabetes mellitus. The present sample consisted of N=459 children from the larger study who had umbilical cord blood plasma assayed for levels of multiple PFAS and who had at least one postnatal magnetic resonance imaging (MRI) scan that met quality control standards. Written informed consent was obtained from participants at recruitment. The GUSTO study was approved by the National Healthcare Group Domain Specific Review Board (Reference #D/09/021) and the SingHealth Centralized Institutional Review Board (Reference #20009/280/D).

### PFAS Estimation

Immediately following delivery, umbilical cord blood was collected in EDTA tubes and centrifuged at 1600 g for 10 min at 4 °C to isolate plasma. Plasma (4 mL) was combined with Trasylol (15 μL) for stabilization and stored at −80 °C in 0.4 mL aliquots until analysis. Ultra performance liquid chromatography-tandem mass spectrometry was used to estimate levels of 12 PFAS, including seven long-chain PFAS: PFOA, PFOS, PFNA, PFDoA, PFHxS, perfluorodecanoic acid (PFDA) and perfluoroundecanoic acid (PFUnDA), and five short-chain PFAS: PFBS, PFBA, perfluoroheptanoic acid (PFHpA), perfluoropentanoic acid (PFPeA), and perfluorohexanoic acid (PFHxA). Detailed information about the quantification of PFAS from cord blood plasma samples has been presented previously (see Chen et al., 2024). PFPeA and PFHxA had low detection rates across samples (0.4–26%) and were excluded from further analysis; PFDoA was detected in 82% of samples but only 9% of samples were above the instrument’s level of quantification (LOQ); therefore, it was excluded from the analysis as well. For the remaining 9 PFAS, samples below the LOQ or limit of detection (LOD) were imputed with LOQ/2 and LOD/2, respectively (Chen et al. 2024). Information about LOQ and LOD for each analyte is reported in the Supplement (Supplemental Table 1). Prior to analysis, concentrations were log_2_ transformed to obtain normal distributions.

### Covariates

We covaried for sociodemographic variables that could confound the associations of PFAS with brain structure, including ethnicity (Chinese, Malay, Indian), and socioeconomic status (SES; Gleason et al., 2025; McAdam & Bell, 2023; Sagiv et al., 2015). SES was operationalized using mother’s report of their monthly household income in Singaporean dollars (1 = $0 – $999, 2 = $1000-$1999, 3 = $2000-$3999, 4 = $4000-$5999, 5 >=$6000) and their highest educational attainment (1 = No education, 2 = Primary, 3 = Secondary or GCE O/N Levels, 4 = Technical or Vocational Training, 5 = GCE A Levels or Polytechnic Diploma, 6 = University Bachelors, Masters or PhD); scores on these variables were z-scored and averaged. In addition, we adjusted for factors that might influence the measurement of brain metrics, including the MRI scanner used, the child’s biological sex and, for models of cortical surface area, thickness, and subcortical volumes, the child’s total intracranial volume.

### MRI Data Acquisition

To minimize motion during MRI scanning, children underwent an MRI home training program prior to the MRI visit and completed on-site training at KK Women’s and Children’s hospital. Structural and diffusion images were acquired in the same session at ages 4.5, 6, 7.5, and 10.5 years using 3T Siemens scanners (Siemens Healthineers, Erlangen, Germany). Scanning at ages 4.5 and 6 years was conducted with a Siemens Magnetom Skyra, and at ages 7.5 and 10.5 years with a Siemens Magnetom Prisma; scanner was included as a fixed effect covariate in the statistical analysis to account for scanner-related differences in brain metrics. High-resolution T1-weighted anatomical volumes were obtained at all timepoints using a magnetization-prepared rapid gradient-echo (MPRAGE) sequence: repetition time = 2000 ms; echo time = 2.08 ms; inversion time = 877 ms; flip angle = 9°; field of view = 192 × 192 mm²; acquisition matrix = 192 × 192; slice thickness = 1.0 mm; total acquisition time was approximately 3.5 minutes. These parameters were held constant across scanner platforms, except number of slices (160 at ages 4.5 and 6 years; 192 at ages 7.5 and 10.5 years). Diffusion-weighted images were acquired using a single-shot echo planar imaging (EPI) sequence: field of view = 192 × 192 mm²; voxel size = 2 mm isotropic; repetition time = 8200 ms; echo time = 85 ms; flip angle = 90°; 30 non-collinear directions; b values = 0, 1000 s/mm²; acceleration factor = 3.

### Cortical and Subcortical Structural Processing

All scans were visually inspected for motion artifacts and poor white matter/gray matter segmentation by a trained neuroradiologist; those that did not pass quality control guidelines were removed prior to further processing (Ducharme et al. 2016). Images were processed with FreeSurfer image analysis suite (Version 7.1.1) using the standard recon-all pipeline (http://surfer.nmr.mgh.harvard.edu/) for anatomical segmentation. Cortical thickness and surface area were derived for each hemisphere using FreeSurfer’s surface-based cortical parcellation (aparc), which parcellates the cortical mantle into 34 regions per hemisphere based on the Desikan-Killiany atlas (Desikan et al. 2006). Subcortical gray matter volumes were derived using FreeSurfer’s automated volumetric segmentation (aseg), which labels subcortical structures based on a probabilistic atlas (Fischl et al. 2002). Bilateral volumes for the thalamus, caudate, putamen, pallidum, hippocampus, amygdala, nucleus accumbens, and brain stem were extracted separately for the left and right hemispheres. Intracranial volume (ICV) was also extracted from the aseg output and used as a covariate in subsequent statistical models.

### White Matter Tract Processing

Diffusion images at each timepoint were preprocessed with FMRIB’s Software Library (FSL, v6.0), except for eddy correction at the 4.5- and 6-year scans, which used an earlier release of FSL’s eddy tool (v5.0.2.2). FMRIB’s eddy tool iteratively predicted and estimated the eddy-current distortions and performed a single resampling of the data incorporating the susceptibility estimates. For images acquired at 7.5 years and 10.5 years, TOPUP correction was applied prior to eddy, wherein b0 volumes were extracted from the diffusion data and concatenated with reverse phase-encoded images to estimate susceptibility-induced distortions. Images were then corrected for eddy currents and susceptibility-by-motion interactions, with outlier slices simultaneously replaced using a Gaussian Process prediction with a threshold of 4 standard deviations. The corrected data from all timepoints were skull-stripped again to isolate brain tissue and denoised using a local principal component analysis (PCA) method (Huang et al. 2025). Average absolute motion across all volumes was computed using FMRIB’s eddy_quad function to assess MRI motion; sensitivity analysis was carried out on a subset of participants with < 3mm movement to demonstrate that the findings were not driven by motion (see Supplement).

Next, within-voxel probability density functions of the principal diffusion direction were estimated using Markov Chain Monte Carlo sampling in FSL’s BEDPOSTX tool. Automated probabilistic tractography was performed using the autoPtx plugin to reconstruct 27 standard white matter pathways in each participant’s native diffusion space, comprising 12 bilateral tracts — the acoustic radiation, anterior thalamic radiation, cingulum (cingulate gyrus and hippocampal portions), corticospinal tract, inferior fronto-occipital fasciculus, inferior longitudinal fasciculus, medial lemniscus, posterior thalamic radiation, superior longitudinal fasciculus, superior thalamic radiation, and uncinate fasciculus — and 3 non-lateralized midline tracts: the forceps minor, forceps major, and middle cerebellar peduncle. Voxel values in each raw tract map were normalized by dividing by the total number of successfully generated streamlines for each pathway, and a strict threshold of 0.001 was then applied to exclude spurious connections and produce a conservative binary mask of each tract’s core. These subject-specific tract masks were overlaid onto the native diffusion maps to extract mean Fractional Anisotropy (FA) for each pathway at each timepoint.

### Analytic Plan

Given the longitudinal measurements of structural brain metrics across childhood, we fit longitudinal mixed effects models (using lme4 in R Version 4.4.3) with participant-level random intercepts to account for repeated measurements within individuals. These models use all available information for estimation, allowing participants to be included in the analysis even if they are missing outcome data at a particular wave (Snijders and Bosker 2011). We first determined whether the developmental trajectory, i.e., age-related change, in each outcome was better characterized by a linear or a quadratic function of age. For each structural metric (e.g., regional cortical thickness and surface areas, subcortical volumes, and FA of white matter tracts), we compared models with a linear age term versus both a linear and quadratic age term using likelihood ratio tests (LRTs). Subsequent analyses used the linear or quadratic modeling of age-related change that was best for each brain metric.

We then tested associations of individual PFAS with each brain metric and with age-related change in brain metrics. To do so, each longitudinal mixed model included a “PFAS x Age” interaction term (and, if applicable, an additional quadratic “PFAS x Age^2^” interaction term). Age was centered at 4.5 years of age (i.e., age at the initial assessment) in order to estimate the effect of doubling of the given PFAS on brain structures when the child was 4.5 years of age. PFAS were centered at their grand means so that age terms reflected age-related change for the overall sample, and interaction terms reflected differences in age-related trajectories per doubling of the respective PFAS. Trajectory modification by PFAS was evaluated by LRTs comparing models with only PFAS main effects to nested models that added the PFAS by age interaction terms.

All estimates were adjusted for the child’s sex, socioeconomic status, ethnicity, and the MRI scanner; cortical structural measures were additionally adjusted for total intracranial volume. The false discovery rate (FDR) was controlled at α =.05 using the Benjamini–Hochberg (BH) procedure. Consistent with prior studies of PFAS and brain development (Haraburda et al. 2026; Shen et al. 2021; Wang et al. 2025; Weng et al. 2020), corrections were performed separately for each PFAS and were conducted within each outcome domain: cortical thickness, cortical surface area, subcortical volumes, and white matter tract FA.

## Results

### Descriptive Statistics

Descriptive statistics for sample demographic characteristics and mean levels of PFAS in cord blood samples are reported in Table 1. To yield the largest sample, children were included in as many assessment timepoints as possible, even if they missed an MRI scan at a prior timepoint or a particular scan was unusable. In total, 459 children were included in the structural MRI sample: 96 (21%) of children had four usable structural MRI scans, 142 (31%) had three usable scans, 109 (24%) had two usable scans, and 112 (24%) had one usable scan (structural MRI: *n*_Y4.5_=187, *n*_Y6_=295, *n*_Y7.5_=342, *n*_Y10.5_=316). For the diffusion scan, 455 children were included in the sample: 109 (24%) of children had four diffusion images, 125 (27%) had three images, 108 (24%) had two images, and 113 (25%) had one image (diffusion MRI: *n*_Y4.5_=245, *n*_Y6_=291, *n*_Y7.5_=311, *n*_Y10.5_=293). Number of scans completed did not differ as a function of PFAS exposure (all p>.05) or the child’s sex (t=-0.264, p=0.792), but did differ by ethnicity (F=5.806, p=.003), with Malay participants completing more scans than both Chinese (p=.038) and Indian (p=.004) participants; in addition, participants with higher SES completed fewer scans (r=-0.110, p=.018).

**Table 1.**
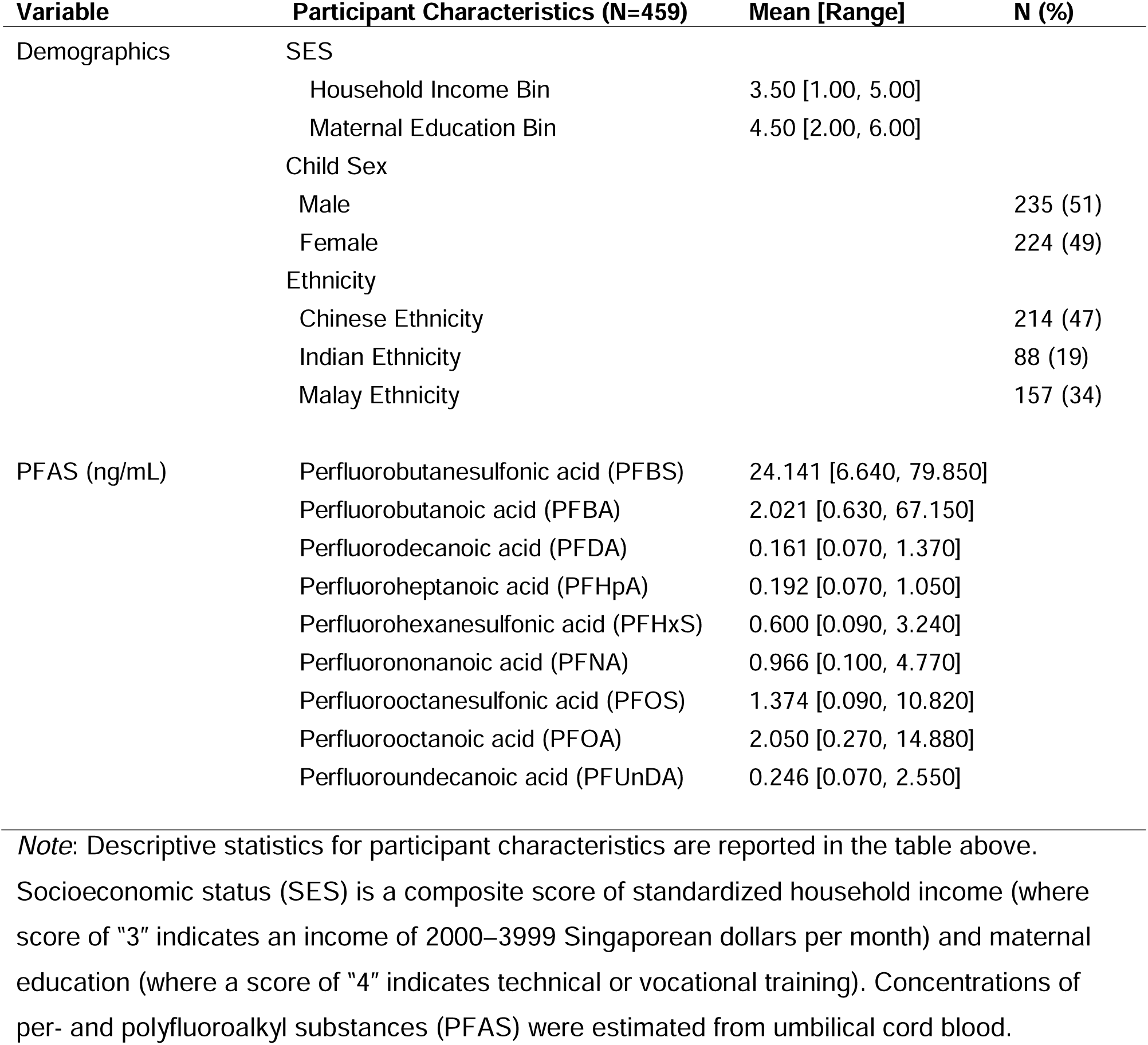
Descriptive characteristics of the sample.

### PFAS Associations with Cortical Structure

We examined associations of prenatal exposure to PFAS with structural development of the cortex across childhood. Specifically, we conducted longitudinal mixed models to estimate the associations of individual PFAS with regional cortical thickness and surface area at 4.5 years of age, as well as the associations of PFAS with developmental trajectories of cortical thickness and surface area across childhood, after accounting for the FDR. We found that three long-chain PFAS (PFDA, PFHxS, and PFNA) were significantly associated with lower surface area of the right paracentral lobule, a region involved in integrating sensorimotor information, at 4.5 years of age. Specifically, a doubling in cord blood levels of PFDA, PFHxS, and PFNA was associated with 49.77 mm^2^, 39.19 mm^2^, and 51.35 mm^2^ lower surface area of the right paracentral lobule, respectively (PFDA: b=-49.765 [-78.502, −21.027], q=.048; PFHxS: b= −39.185 [-61.389, −16.981], q=.038; PFNA: b=-51.346 [-78.277, −24.414], q=.013; see Figure 1 and Supplemental Table 2). No other PFAS (i.e., PFOA, PFOS, PFUnDA, PFHpA, PFBS, PFBA) were significantly associated with thickness or surface area of regional cortical structures at 4.5 years of age after FDR corrections.

**Figure 1.**
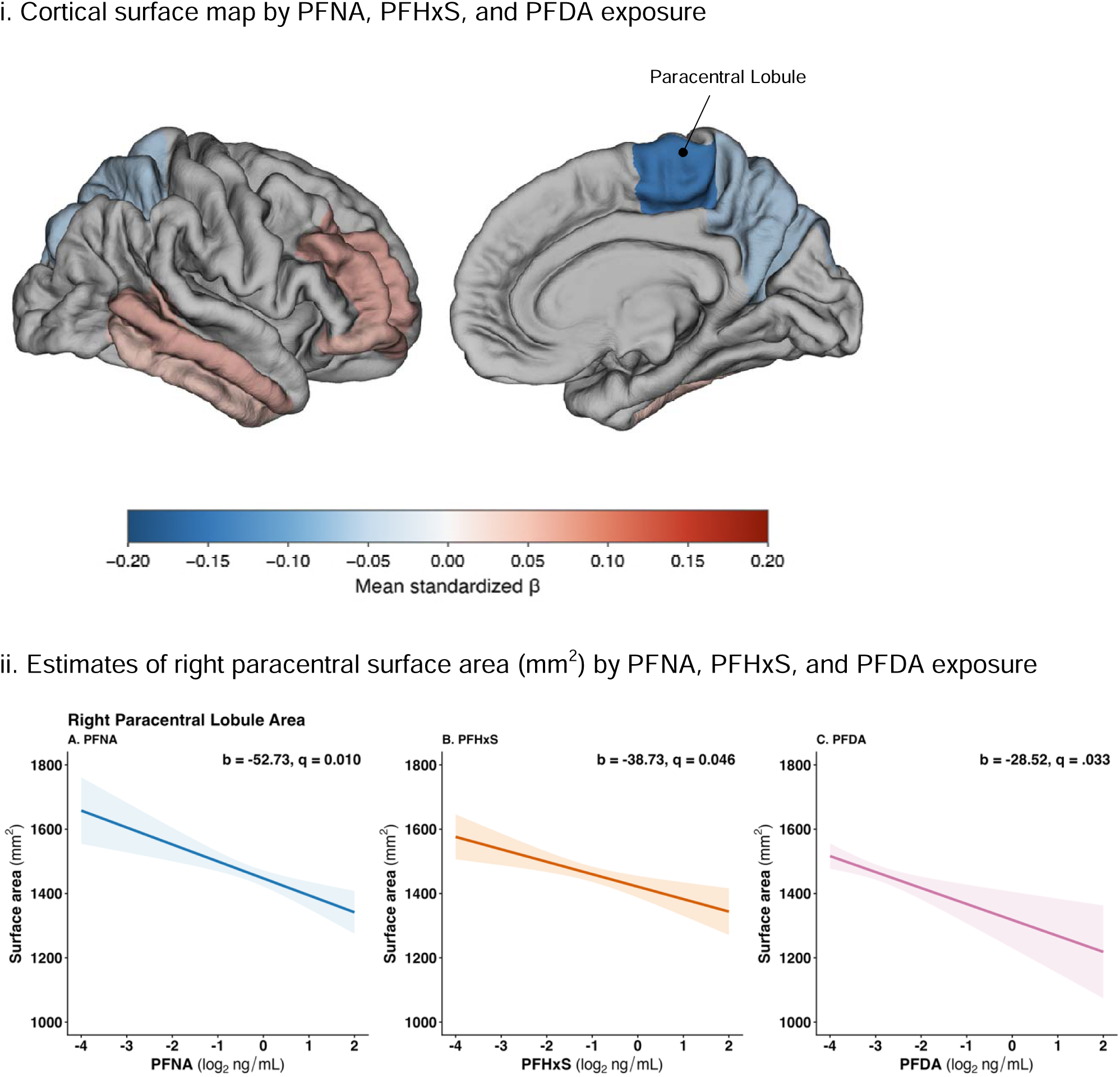
Associations of PFNA, PFHxS, and PFDA with right-hemisphere cortical surface area at 4.5 years of age *Note:* The top panel (i) displays the right-hemisphere pial cortical surface (fsaverage template), showing the mean standardized β (centered at age 4.5 years) for each Desikan-Killiany region. Color encodes both the sign and magnitude of the effect (blue = negative β, smaller surface area with higher prenatal PFAS exposure; red = positive β), while opacity encodes statistical tier: regions where at least one PFAS chemical reached FDR significance (*q* < 0.05) are shown at full opacity, regions where the strongest effect reached only nominal significance (uncorrected *p* < 0.05) are shown at reduced opacity, and regions with no association at *p* < 0.05 are unfilled (gray). All three FDR-significant surface-area effects (PFNA, PFHxS, and PFDA) converged on the right paracentral lobule; the left hemisphere is omitted. For visualization, the plot displays associations averaged across PFNA, PFHxS and PFDA, however the compounds are evaluated individually in statistical models. The bottom panel (ii) displays predicted values of surface area of the right paracentral lobule at 4.5 years of age by prenatal exposure to PFNA, PFHxS, and PFDA from linear mixed effect models. Unstandardized beta estimates reflect the association of a doubling in the respective PFAS with surface area (mm^2^) after adjusting for socioeconomic status, ethnicity, sex, intracranial volume, and MRI scanner. Adjusted p-values (q) accounted for FDR using the Benjamini-Hochberg correction.

Next, we tested whether prenatal exposure to PFAS was associated with alterations in the longitudinal trajectory of thickness and surface area in cortical structures across childhood. Although we had found significant associations of PFDA, PFHxS, and PFNA with reduced surface area of the paracentral lobule at 4.5 years of age, none was associated with alterations in the trajectory of surface area growth of the paracentral lobule across childhood after FDR corrections; that is, model fit was not significantly improved when including PFAS interactions with linear and quadratic age terms (PFDA: χ²=6.140, q=.650; PFHxS: χ²=8.351, q=.289; PFNA: χ²=10.691, q=.278; see Supplemental Table 2). Together, our findings suggest that children who had higher prenatal exposure to these long-chain PFAS had lower baseline surface area of the paracentral lobule, but their trajectory of surface area growth across childhood did not differ significantly from that of children with lower PFAS exposure.

There were no other associations of long-chain PFAS (i.e., PFOA, PFOS, PFUnDA) with longitudinal trajectories of regional cortical surface area and thickness. However, we did find significant associations of the short-chain PFHpA with altered trajectories of surface area growth in two frontal regions. First, PFHpA was associated with an altered trajectory of surface area growth in the left rostral anterior cingulate cortex (rACC; χ²=11.405, q=.025), a limbic region involved in emotion regulation. Although PFHpA was not associated with surface area in the left rACC at 4.5 years of age (b=0.042 [-9.127, 9.212], q=.993), children with higher PFHpA had significantly steeper (i.e., faster) surface area growth in the left rACC across childhood (b=1.896 [0.799, 2.994], q=.023; see Figure 2 and Supplemental Table 2). Second, PFHpA was associated with alterations in the trajectory of surface area growth in the right caudal middle frontal gyrus (χ²=14.513, q=.025), a frontal brain region involved in attention and executive control. That is, children with higher PFHpA exposure showed declines in surface area followed by a deceleration of this decline with age (linear age term: b=-31.473 [-47.946, −15.001], q=.013; quadratic age term: b=4.170 [1.756, 6.585], q=.036), reflecting less surface area growth in this region from 4.5 to 10.5 years relative to children with lower PFHpA exposure (see Figure 2 and Supplemental Table 2). Neither of the remaining short-chain PFAS (PFBS, PFBA) was associated with longitudinal trajectories of cortical thickness and surface area across childhood.

**Figure 2.**
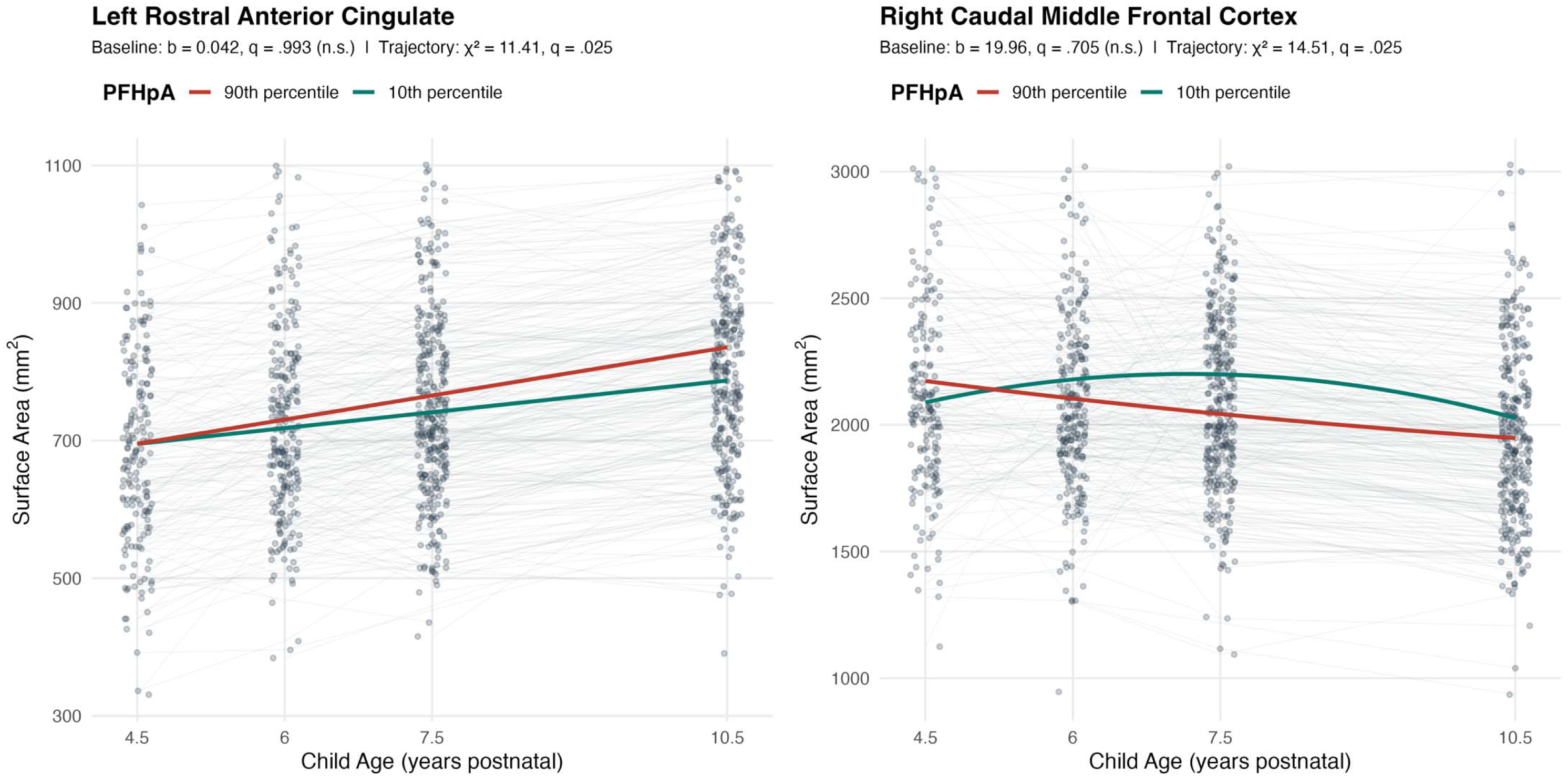
PFHpA associations with longitudinal surface area trajectories of cortical structures across childhood. *Note:* The plots display surface area of the left rostral anterior cingulate and right caudal middle frontal gyrus across childhood. Gray dots denote children’s surface area values after residualizing out model control variables (socioeconomic status, ethnicity, sex, intracranial volume and MRI scanner), and dots are connected across study timepoints with gray lines. PFAS were evaluated as continuous variables in all models, however, for purposes of visualization, the plots display estimated longitudinal trajectories of surface area for a child whose cord blood concentration of PFHpA was at the 90^th^ percentile (red line), as compared to a child whose cord blood concentration of PFHpA was at the 10^th^ percentile (green line). Datapoints have been jittered along the x-axis to enhance visualization.

### Subcortical Volumes

No associations of PFAS with volumes of subcortical structures at 4.5 years of age survived FDR corrections. Similarly, no PFAS were significantly associated with longitudinal trajectories of subcortical volumes from ages 4.5 to 10.5 years of age.

### White Matter Tract Fractional Anisotropy

We examined associations of prenatal exposure to PFAS with FA of key white matter tracts at 4.5 years of age and associations with the developmental trajectory of FA in these tracts across childhood. Of the nine PFAS we examined here, the short-chain PFBS was associated most consistently with FA of white matter tracts at 4.5 years of age, showing significant associations with higher FA in 17 of the 27 white matter tracts examined. Specifically, at age 4.5 years PFBS was associated with higher FA in the bilateral corticospinal tract, the primary pathway for motor information (left hemisphere: b=0.012 [0.007, 0.018], q<.001; right hemisphere: b=0.015 [0.009, 0.021], q<.001; see Figure 3 and Supplemental Table 3). PFBS was also associated with higher FA in six thalamocortical projection tracts that bridge the thalamus to regions of the cortex: the bilateral anterior thalamic radiation (left hemisphere: b=0.007 [0.003, 0.011], q=.007; right hemisphere: b=0.006 [0.002, 0.010], q=.009), the bilateral superior thalamic radiation (left hemisphere: b=0.006 [0.002, 0.011], q=.009; right hemisphere: b=0.006 [0.002, 0.010], q=.012), the left posterior thalamic radiation (b=0.007 [0.001, 0.012], q=.022), and the right acoustic radiation (b=0.007 [0.003, 0.012], q=.007; see Figure 3 and Supplemental Table 3). Finally, PFBS was associated with higher FA in multiple intrahemispheric, long-range association tracts and commissural pathways, including the left inferior fronto-occipital fasciculus (b=0.007 [0.002, 0.011], q=.009), the right inferior fronto-occipital fasciculus (b=0.007 [0.002, 0.013], q=.022), the left inferior longitudinal fasciculus (b=0.006 [0.002, 0.011], q=.011), the left superior longitudinal fasciculus (b=0.005 [0.001, 0.008], q=.022), the left uncinate fasciculus (b=0.004 [0.001, 0.008], q=.041), the right cingulate gyrus segment of the cingulum bundle (b=0.006 [0.001, 0.012], q=.038), the forceps major (b=0.009 [0.004, 0.014], q=.005), the forceps minor (b=0.009 [0.003, 0.014], q=.007), and the middle cerebellar peduncle (b=0.011 [0.004, 0.018], q=.007; see Figure 3 and Supplemental Table 3).

**Figure 3.**
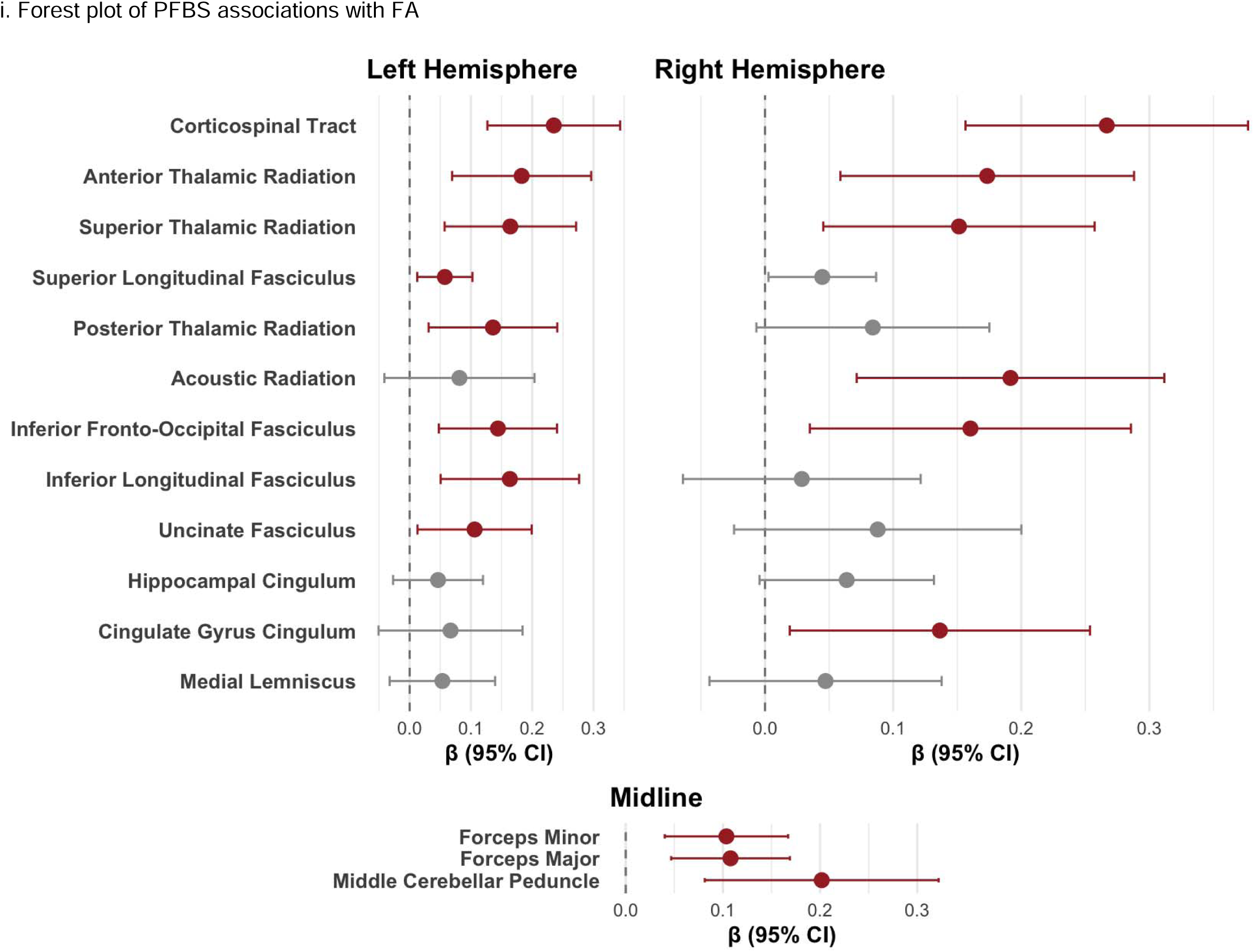
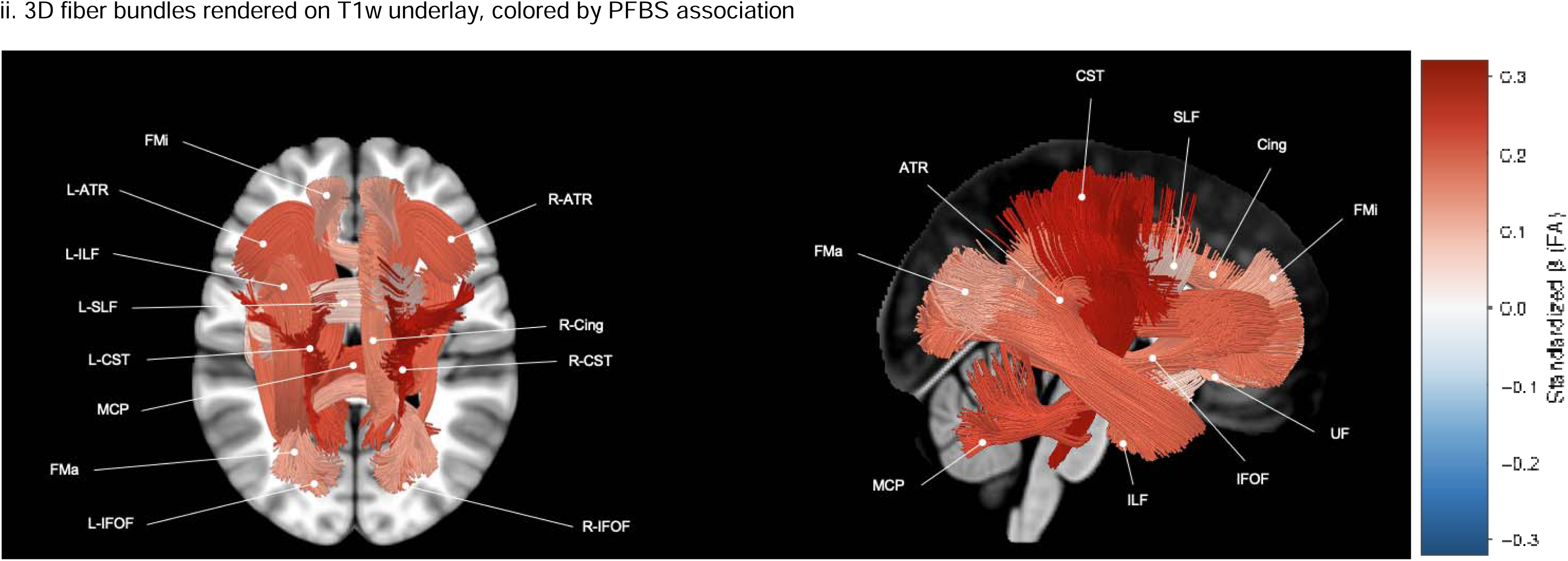
Associations of PFBS with fractional anisotropy of major white matter tracts at 4.5 years of age *Note:* The top panel (i) reports standardized betas (β) and 95% confidence intervals from mixed effect models for the associations of PFBS with FA of individual white matter tracts at 4.5 years of age. Estimates and 95% confidence intervals of tracts that were significantly associated with PFBS after accounting for the false discovery rate are colored in red and insignificant associations are colored in gray. All estimates are adjusted for socioeconomic status, sex, ethnicity, and MRI scanner. The bottom panel displays FDR-significant tract-level associations rendered as 3D fiber bundles on an MNI152 T1-weighted underlay (mid-sagittal slice for the lateral view; axial slice at z = +15 mm for the dorsal view), colored by the mean standardized β (centered at age 4.5 years) for PFBS. Color encodes both the sign and magnitude of the effect (blue = negative β, lower FA with higher prenatal PFAS exposure; red = positive β); only tracts reaching FDR significance (*q* < 0.05) are shown. The panel shows a right-lateral and a dorsal (top-down) view. Streamlines are drawn from the HCP1065 atlas (corticospinal tract, cingulum, inferior fronto-occipital fasciculus, uncinate fasciculus, middle cerebellar peduncle, forceps major/minor); fiber-like curves for tracts available only as probability maps (anterior thalamic radiation, inferior longitudinal fasciculus, superior longitudinal fasciculus, parahippocampal cingulum) were synthesized from the AFQ atlas. Four additional FDR-significant tracts are not rendered because no atlas streamline or probability map was available (right acoustic radiatio and right superior thalamic radiation, left posterior thalamic radiation).

In addition, we examined associations of PFBS with the trajectory of FA in major white matter tracts across childhood. We found that PFBS was associated significantly with alterations in the longitudinal trajectories of FA in 13 white matter tracts: the bilateral corticospinal tract (left hemisphere: χ²=16.337, q=.004; right hemisphere: χ²=16.295, q=.004), the right acoustic radiation (χ²=7.539, q=.048), the bilateral anterior thalamic radiation (left hemisphere: χ²=9.321, q=.033; right hemisphere: χ²=10.065, q=.033), the left superior thalamic radiation (χ²=9.274, q=.033), the left inferior fronto-occipital fasciculus (χ²=7.118, q=.033), the left inferior longitudinal fasciculus (χ²=7.869, q=.044), the left uncinate fasciculus (χ²=8.255, q=.027), the right cingulate gyrus segment of the cingulum bundle (χ²=8.484, q=.039), the forceps major (χ²=8.737, q=.038), the forceps minor (χ²=11.465, q=.027), and the middle cerebellar peduncle (χ²=8.273, q=.039; see Supplemental Table 3). As visualized in Figure 4, children primarily demonstrated a quadratic pattern of FA increase in these tracts across childhood; compared to children with lower prenatal PFBS exposure, those with higher exposure had steeper initial declines in FA followed by subsequent increases, such that they were more similar to youth with lower exposure by 10.5 years of age—a pattern that could indicate possible convergence to normative levels with age.

**Figure 4.**
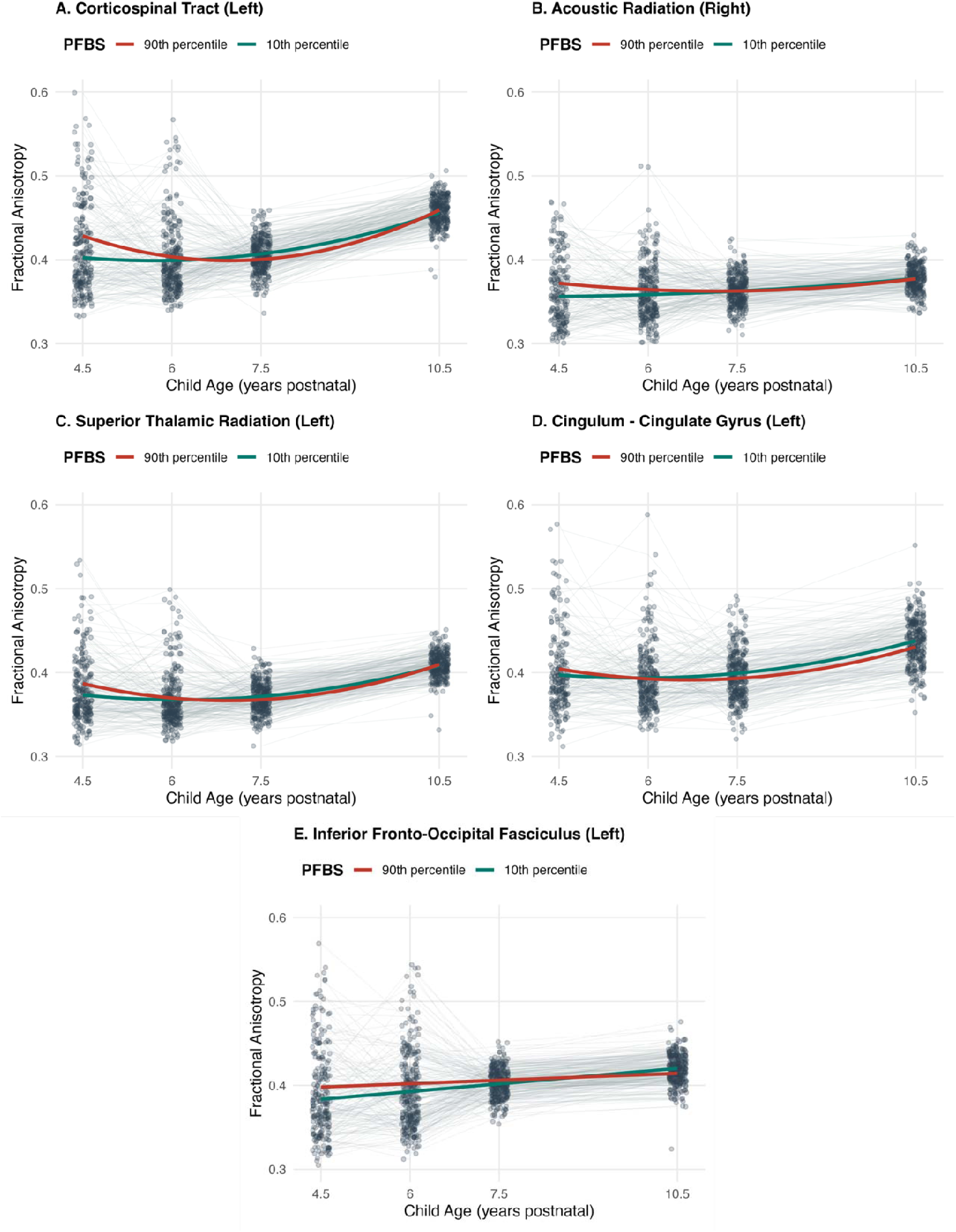
Selected PFBS associations with developmental trajectories of fractional anisotropy in white matter tracts across childhood. *Note:* The plots display fractional anisotropy (FA) across childhood of five white matter tracts that had both significant baseline and longitudinal trajectory associations with PFBS, which included A) the left corticospinal tract, B) the right acoustic radiation, C) the left superior thalamic radiation, D) the cingulate gyrus portion of the cingulum and E) the left inferior fronto-occipital fasciculus. Gray dots denote children’s FA values after residualizing out model control covariates (i.e., socioeconomic status, ethnicity, sex and MRI scanner), and dots are connected across study timepoints with gray lines. PFAS were evaluated as continuous variables in all models, however, for purposes of visualization, the plots display estimated longitudinal trajectories of FA in each white matter tract for a child whose cord blood concentration of PFBS was at the 90^th^ percentile (red line), as compared to a child whose cord blood concentration of PFBS was at the 10^th^ percentile (green line). Datapoints have been jittered along the x-axis to enhance visualization.

In contrast to findings obtained for PFBS, higher cord blood concentration of the long-chain legacy compound, PFOA, was associated with *lower* FA in six white matter tracts at 4.5 years of age: the left cingulate gyrus segment of the cingulum bundle (b=-0.007 [-0.012, −0.002], q=.018), the right hippocampal segment of the cingulum bundle (b=-0.005 [-0.008, −0.002], q=.006), the right inferior fronto-occipital fasciculus (b=-0.006 [-0.011, −0.001], q=.047), the middle cerebellar peduncle (b=−0.009 [−0.014, −0.003], q=.012), the forceps major (b=−0.009 [−0.013, −0.005], q<.001), and the forceps minor (b=−0.008 [−0.012, −0.004], q=.005; see Supplemental Table 3). In addition, PFOA was associated with altered longitudinal trajectories of FA in seven white matter tracts: the left cingulate gyrus segment of the cingulum bundle (χ²=13.875, q=.007), the right hippocampal segment of the cingulum bundle (χ²=15.864, q=.003), the right inferior fronto-occipital fasciculus (χ²=10.219, q=.025), the bilateral uncinate fasciculus (left hemisphere: χ²=8.957, q=.015; right hemisphere: χ²=10.084, q=.025), the forceps major (χ²=22.041, q<.001), and the forceps minor (χ²=15.815, q=.003; see Supplemental Table 3). In each of these tracts, the negative association of PFOA with FA at 4.5 years of age was attenuated with age (i.e., positive PFAS x age interaction terms; see Supplemental Table 3), such that children with higher PFOA exposure showed a pattern of increasing FA across childhood relative to those with lower exposure. No other long or short-chain PFAS were associated with FA at age 4.5 years or with the trajectories of FA in major white matter tracts across childhood.

## Discussion

This multimodal imaging study was designed to examine the associations of prenatal exposure to PFAS with brain development across childhood. Higher prenatal exposure to specific PFAS was associated with alterations during childhood in surface area of frontal and paracentral cortical structures related to sensorimotor and limbic functioning and with FA in key white matter tracts. At 4.5 years of age, associations with surface area were most consistent for long-chain compounds, including PFNA, PFDA and PFHxS. These associations were localized to lower surface area of the right paracentral lobule rather than reflecting widespread cortical effects. As children aged from 4.5 to 10.5 years, PFHpA was associated with faster surface area growth in the left rACC, a region that serves emotion regulatory functions, and with less growth in the right caudal middle frontal gyrus, a region implicated in attention and executive functioning. Contrary to our hypothesis, however, levels of PFAS in cord blood were not associated with altered trajectories of cortical thickness or subcortical volumes across childhood. Finally, prenatal exposure to PFBS, a newer, short-chain PFAS, was associated with widespread alterations in white matter fiber integrity, with higher PFBS linked to higher FA across major long-range projection and inter- and intrahemispheric association tracts. Conversely, higher levels of the legacy compound PFOA were associated with lower FA in several of these tracts. These white matter associations were most pronounced at 4.5 years of age, with evidence of attenuation in several tracts as children aged. Given the increasing production of PFBS, which was detected at high levels in our sample, these findings raise concerns about the impact of short-chain replacement PFAS on children’s neurodevelopment.

The most consistent findings of prenatal PFAS on brain development involved associations between higher PFBS concentrations and greater FA across the majority of white matter tracts assessed at 4.5 years of age. Although FA can reflect greater fiber bundle density and/or myelination, reduced fiber branching and crossing can also lead to higher FA, which has been linked to risk for neurodevelopmental disorders such as ADHD (Davenport et al. 2010). FA generally increases across childhood with ongoing myelination and fiber packing (Lebel et al. 2012); thus, earlier organization of white matter tracts could indicate a deviation from normative neurodevelopment. Prior studies of the effects of prenatal PFAS exposure on white matter microstructure have yielded mixed results. For example, Barron et al. (2025) found that higher prenatal PFOA and PFNA were associated with greater integrity of the corpus callosum at five years of age; however, England-Mason et al. (2025) reported that PFOS and PFHxS were associated with *lower* FA in the corpus callosum. In the present study, we found that PFAS had different associations with the forceps major and minor of the corpus callosum: specifically, whereas higher levels of PFBS were related to *greater* FA in these tracts, higher levels of PFOA were related to *lower* FA. This directional divergence might reflect the ongoing industrial replacement of the legacy compound PFOA with short-chain compounds such as PFBS (Kannan et al. 2026). This same divergence for PFBS and PFOA was also apparent in the middle cerebellar peduncle, which has been documented in other developmental cohorts. For example, Wang et al. (2025) found that PFDoA, a long-chain PFAS, was negatively associated with FA and related diffusion metrics in the inferior and superior cerebellar peduncles among male adolescents, and Shen et al. (2021) found smaller cerebellar volume in relation to several long-chain PFAS. Together, these findings suggest that exposure to PFAS during gestation plays a role in the development of white matter microstructure; however, given the limited investigation of the effects of newer, replacement PFAS, the direction of these associations remains unclear.

The mechanisms that might explain how PFAS influences white matter microstructure are also not well understood. One possibility is that exposure to PFAS during gestation disrupts important myelination processes. For instance, PFAS have been shown to activate peroxisome proliferator-activated receptors (PPARs; Evans et al. 2022) and disrupt thyroid hormone signaling (Coperchini et al. 2021), both of which regulate oligodendrocyte development and the expression of myelin-related genes (Calzà, Fernández, and Giardino 2015; Han et al. 2025). PFAS have also been found to induce oxidative stress (Taibl et al. 2022), and developing oligodendrocytes are particularly vulnerable to oxidative injury (Ljubisavljevic 2016). We found that the strongest associations of PFBS exposure were with the corticospinal tract, the main motor projection pathway. This finding could reflect the developmental sequence of white matter maturation, in which sensorimotor projection tracts myelinate earlier than do frontal association tracts (Dubois et al. 2014). The attenuation of these associations with age may indicate a developmental “catch-up” in tract maturation as children aged or, alternatively, could be a consequence of changing environmental exposures following birth. Further longitudinal research measuring exposure to PFAS and other environmental contaminants multiple times across childhood is needed to examine these possibilities.

Our findings concerning cortical surface area extend results of previous studies. We found significantly reduced surface area of the right paracentral lobule at 4.5 years as a function of prenatal exposure to PFNA, PFDA and PFHxS. The paracentral lobule sends motor output via the corticospinal tract and receives somatosensory input via the superior thalamic radiation, both of which showed altered development with exposure to PFAS. Together, these findings may implicate PFAS in the development of sensorimotor circuits and help to explain prior associations between PFAS exposure and children’s motor delays (Luo et al. 2022). Our results also highlight the importance of developmental timing: although PFHpA levels were not associated with left rACC surface area at 4.5 years of age, youth with higher PFHpA levels had faster surface area growth in the left rACC through age 10.5 years, a finding consistent with Wang et al.’s (2025) linking of prenatal exposure to PFDA to greater surface area of the left rACC in 14- to 15-year-old adolescents. Atypical surface area expansion of the rACC combined with slower growth of the caudal middle frontal gyrus, which supports executive control (Andersson et al. 2009), could influence children’s emotion regulatory skills (Etkin, Egner, and Kalisch 2011). Considering this, our findings may provide context for the documented association of PFAS with behavioral problems (Kim et al. 2023; Tillaut et al. 2023). Finally, contrary to our hypothesis, PFAS were not associated with subcortical volumes despite their presumed higher PFAS concentrations. However, Weng et al. (2020) reported links between PFAS and reduced resting-state functional activity in the putamen, caudate, insula, and pallidum—suggesting that functional neural activity might be more sensitive to PFAS exposure than is structural volume.

We should note four limitations of this study. First, prenatal exposure to PFAS was estimated from concentrations in cord blood; therefore, we are not able to examine the influence of later PFAS exposure on brain development. Future research examining PFAS exposure across childhood could elucidate the impact of continuing PFAS burden on the developing brain and clarify the effects of exposure on brain structure during sensitive periods of development. Second, we examined structural metrics of brain development, including surface area, thickness, and volume, as well as diffusional properties of major white matter tracts. We did not assess whether functional properties of these regions are affected by exposure to PFAS, a question that should be examined in future research. Third, we did not obtain information regarding possible confounding variables such as maternal diet. For example, certain food products, like seafood, can both heighten PFAS exposure and be an important source of omega-3 fatty acids that support fetal brain development (Liu et al. 2017). Finally, the data analyzed in this study were obtained from mother-child dyads in Singapore, where PFBS levels are high (Chen et al. 2017). Thus, our results may not generalize to populations with differing exposure to PFAS, and our findings should be replicated in geographically and demographically diverse cohorts.

Despite these limitations, this is the largest multimodal neuroimaging study of prenatal PFAS exposure and children’s brain development, and the only investigation to evaluate these associations longitudinally across childhood. Further, this is the only neuroimaging study to examine the effects of newer, emerging PFAS on children’s brain development. We demonstrated in this study that early-life PFAS exposure is associated with alterations in brain development across childhood, most consistently in white matter microstructure. It will be important in future research to examine long-term consequences of exposure to PFAS, particularly of exposure to newer replacement PFAS, on both structural and functional metrics of children’s brain development.

## Notes

https://gustodatavault.sg/

### Competing Interest Statement

The authors have declared no competing interest.

### Funding

- National Institute of Mental Health: R37MH101495 to IHG, F32MH135657 to JPU
- Brain & Behavior Research Foundation: Young Investigator Grant #31350 to JLB
- Stanford Maternal and Child Health Institute: Transdisciplinary Initiatives Program award to IHG
- Stanford Center on Longevity: New Map of Life Fellowship to SR
- Singapore Ministry of Health: MOH-001273-00 to APT

