## Supplementary Material for "Prenatal exposure to PFAS and multimodal structural brain development across childhood"

**Supplemental Table 1.** Levels and detection of PFAS in cord blood

| **Compound** | **N** | **Mean [Min, Max]** | **IQR** | **LOD** | **N (%)**  **< LOD** | **LOQ** | **N (%)**  **< LOQ** |
| --- | --- | --- | --- | --- | --- | --- | --- |
| *Analyzed* |  |  |  |  |  |  |  |
| Perfluorobutanesulfonic acid (PFBS) | 459 | 24.141 [6.640, 79.850] | 15.795 | 0.052 | 0 (0.0%) | 3.333 | 0 (0.0%) |
| Perfluorobutanoic acid (PFBA) | 459 | 2.021 [0.630, 67.150] | 0.800 | 0.273 | 6 (1.3%) | 0.333 | 6 (1.3%) |
| Perfluorodecanoic acid (PFDA) | 459 | 0.161 [0.070, 1.370] | 0.080 | 0.007 | 0 (0.0%) | 0.067 | 20 (4.4%) |
| Perfluoroheptanoic acid (PFHpA) | 459 | 0.192 [0.070, 1.050] | 0.120 | 0.033 | 83 (18.1%) | 0.067 | 137 (29.8%) |
| Perfluorohexanesulfonic acid (PFHxS) | 459 | 0.600 [0.090, 3.240] | 0.400 | 0.016 | 1 (0.2%) | 0.067 | 5 (1.1%) |
| Perfluorononanoic acid (PFNA) | 459 | 0.966 [0.100, 4.770] | 0.665 | 0.011 | 0 (0.0%) | 0.067 | 0 (0.0%) |
| Perfluorooctanesulfonic acid (PFOS) | 459 | 1.374 [0.090, 10.820] | 0.830 | 0.018 | 0 (0.0%) | 0.067 | 0 (0.0%) |
| Perfluorooctanoic acid (PFOA) | 459 | 2.050 [0.270, 14.880] | 1.480 | 0.006 | 0 (0.0%) | 0.067 | 0 (0.0%) |
| Perfluoroundecanoic acid (PFUnDA) | 459 | 0.246 [0.070, 2.550] | 0.130 | 0.007 | 2 (0.4%) | 0.067 | 14 (3.1%) |
| *Excluded* | **N** | **Mean [Min, Max]** | **IQR** | **LOD** | **N (%)**  **< LOD** | **LOQ** | **N (%)**  **< LOQ** |
| Perfluorododecanoic acid (PFDoA) | 459 | 0.101 [0.070, 0.290] | 0.050 | 0.009 | 85 (18.5%) | 0.067 | 416 (90.6%) |
| Perfluorohexanoic acid (PFHxA) | 459 | 0.670 [0.490, 0.850] | 0.180 | 0.080 | 457 (99.6%) | 0.333 | 457 (99.6%) |
| Perfluoropentanoic acid (PFPeA) | 459 | 0.162 [0.070, 0.450] | 0.080 | 0.067 | 342 (74.5%) | 0.067 | 342 (74.5%) |

*Note:* Concentrations are in ng/mL. IQR = interquartile range. LOD= limit of detection; LOQ = limit of quantification.

**Supplemental Figure 1.** Bivariate correlations of levels of individual PFAS in cord blood plasma
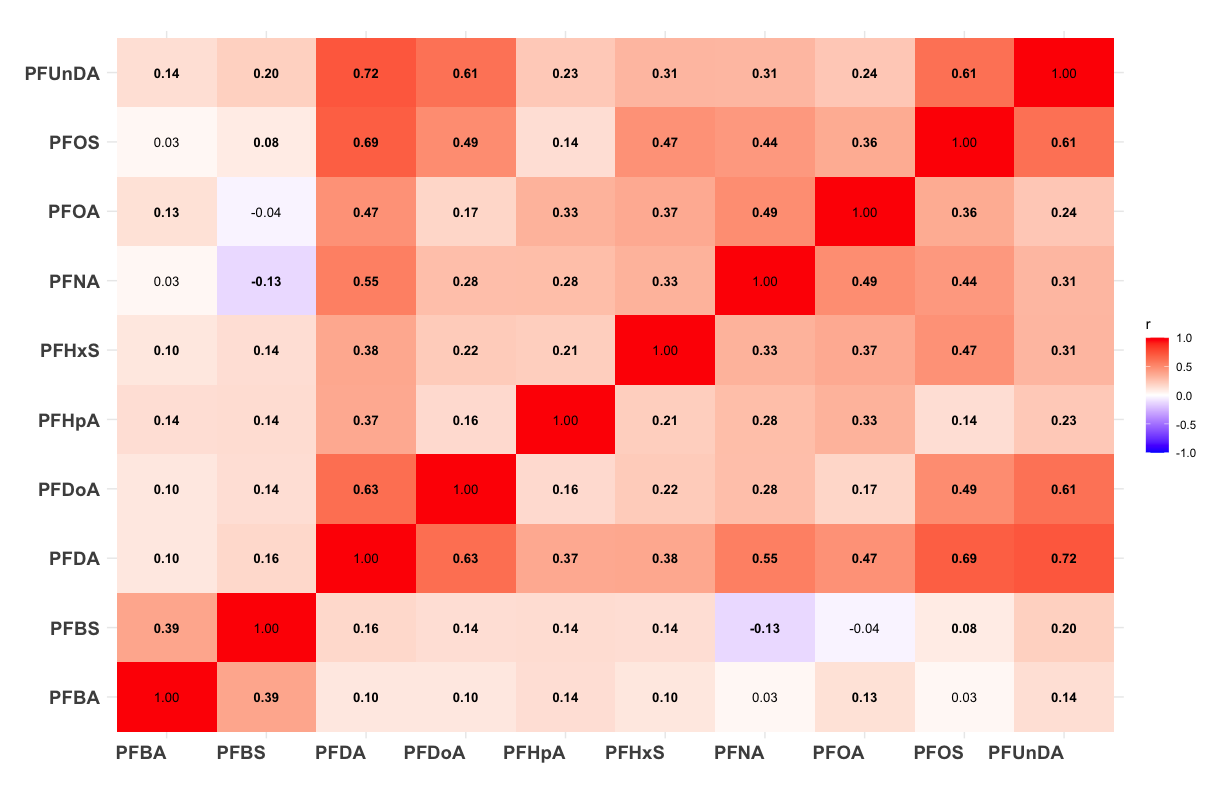

*Note:* The plot reports bivariate correlations between all PFAS detected in cord blood samples. Significant correlations are bolded.

**Supplemental Table 2.** Significant PFAS associations with baseline (4.5 years of age) and developmental trajectories of cortical surface area across childhood

|  |  | **PFAS** |  | **PFAS x Age** |  | **PFAS x Age²** |  | **Trajectory** |  |
| --- | --- | --- | --- | --- | --- | --- | --- | --- | --- |
| **Structure** | **PFAS** | **B [95% CI]** | **p (q)** | **B [95% CI]** | **p (q)** | **B [95% CI]** | **p (q)** | **LRT χ²** | **p (q)** |
| Paracentral Lobule (R) | PFDA | **-49.765 [-78.502, -21.027]** | .001 (.048) | 18.608 [2.177, 35.038] | .026 (.360) | -2.217 [-4.617, 0.183] | .070 (.491) | 6.14 | .046 (.650) |
| Paracentral Lobule (R) | PFHxS | **-39.185 [-61.389, -16.981]** | .001 (.038) | 16.670 [3.610, 29.729] | .012 (.211) | -1.945 [-3.873, -0.017] | .048 (.517) | 8.351 | .015 (.289) |
| Paracentral Lobule (R) | PFNA | **-51.346 [-78.277, -24.414]** | <.001 (.013) | 24.749 [9.071, 40.427] | .002 (.137) | -3.128 [-5.444, -0.813] | .008 (.400) | 10.691 | .005 (.278) |
| Caudal Middle Frontal Gyrus (R) | PFHpA | 19.960 [-9.767, 49.686] | .188 (.705) | -31.473 [-47.946, -15.001] | <.001 (.013) | 4.170 [1.756, 6.585] | .001 (.036) | **14.513** | .001 (.025) |
| Rostral Anterior Cingulate Cortex (L) | PFHpA | 0.042 [-9.127, 9.212] | .993 (.993) | 1.896 [0.799, 2.994] | .001 (.023) | - | - | **11.405** | .001 (.025) |

*Note:* Significant FDR-corrected associations of PFAS with surface area of regional cortical structures from longitudinal mixed models are reported above. The “PFAS B [95% CI]” term reflects the estimated effect and 95% confidence interval of a doubling in the given PFAS on surface area at 4.5 years postnatal (i.e., the initial assessment). Uncorrected p-values (p) and FDR-corrected p-values (q) are reported for each estimate. Each model additionally included either a linear “PFAS x Age” interaction term or both a linear “PFAS x Age” and quadratic “PFAS x Age^2^” interaction term depending on whether modeling fit was improved when allowing for a nonlinear age trend. Models that were better fit linearly report dashes in the nonlinear columns. The significance of PFAS associations with age trajectories was determined from the likelihood ratio test (LRT) comparing model fit with and without the interaction of PFAS with linear and (if applicable) quadratic age terms. Bolded text indicates significant PFAS associations at baseline (Beta [PFAS]) or significant PFAS associations with age trajectories (LRT χ²) after FDR corrections.

**Supplemental Table 3**. Significant PFAS associations with FA in white matter tracts at 4.5 years postnatal and links with age-trends

|  |  | **PFAS** |  | **PFAS x Age** |  | **PFAS x Age²** |  | **Trajectory** |  |
| --- | --- | --- | --- | --- | --- | --- | --- | --- | --- |
| **Tract** | **PFAS** | **B [95% CI]** | **p (q)** | **B [95% CI]** | **p (q)** | **B [95% CI]** | **p (q)** | **LRT χ²** | **p (q)** |
| Acoustic Radiation (R) | PFBS | **0.007 [0.003, 0.012]** | .002 (.007) | -0.004 [-0.007, -0.000] | .034 (.060) | <0.001 [-0.000, 0.001] | .116 (.165) | **7.539** | .023 (.048) |
| Anterior Thalamic Radiation (L) | PFBS | **0.007 [0.003, 0.011]** | .002 (.007) | -0.003 [-0.007, -0.000] | .031 (.059) | <0.001 [-0.000, 0.001] | .137 (.183) | **9.321** | .009 (.033) |
| Anterior Thalamic Radiation (R) | PFBS | **0.006 [0.002, 0.010]** | .003 (.009) | -0.004 [-0.007, -0.001] | .005 (.025) | 0.001 [0.000, 0.001] | .020 (.075) | **10.065** | .007 (.033) |
| Cingulum – Cingulate Gyrus (L) | PFOA | **-0.007 [-0.012, -0.002]** | .003 (.018) | 0.005 [0.001, 0.009] | .006 (.028) | -0.001 [-0.001, 0.000] | .056 (.140) | **13.875** | .001 (.007) |
| Cingulum – Cingulate Gyrus (R) | PFBS | **0.006 [0.001, 0.012]** | .022 (.038) | -0.005 [-0.009, -0.001] | .007 (.026) | 0.001 [0.000, 0.001] | .023 (.075) | **8.484** | .014 (.039) |
| Cingulum – Hippocampus (R) | PFOA | **-0.005 [-0.008, -0.002]** | .001 (.006) | 0.005 [0.002, 0.007] | <.001 (.002) | -0.001 [-0.001, -0.000] | <.001 (.003) | **15.864** | <.001 (.003) |
| Corticospinal Tract (L) | PFBS | **0.012 [0.007, 0.018]** | <.001 (<.001) | -0.008 [-0.013, -0.004] | <.001 (.003) | 0.001 [0.000, 0.002] | .001 (.015) | **16.337** | <.001 (.004) |
| Corticospinal Tract (R) | PFBS | **0.015 [0.009, 0.021]** | <.001 (<.001) | -0.008 [-0.013, -0.004] | <.001 (.006) | 0.001 [0.000, 0.002] | .005 (.051) | **16.295** | <.001 (.004) |
| Inferior Fronto-Occipital Fasciculus (L) | PFBS | **0.007 [0.002, 0.011]** | .003 (.009) | -0.002 [-0.003, -0.000] | .008 (.026) | - | - | **7.118** | .008 (.033) |
| Inferior Fronto-Occipital Fasciculus (R) | PFOA | **-0.006 [-0.011, -0.001]** | .010 (.047) | 0.005 [0.002, 0.009] | .005 (.028) | -0.001 [-0.001, -0.000] | .026 (.102) | **10.219** | .006 (.025) |
| Inferior Fronto-Occipital Fasciculus (R) | PFBS | **0.007 [0.002, 0.013]** | .012 (.022) | -0.004 [-0.008, 0.000] | .072 (.108) | <0.001 [-0.000, 0.001] | .157 (.196) | 4.614 | .100 (.158) |
| Inferior Longitudinal Fasciculus (L) | PFBS | **0.006 [0.002, 0.011]** | .005 (.011) | -0.004 [-0.007, -0.001] | .010 (.029) | 0.001 [0.000, 0.001] | .028 (.080) | **7.869** | .020 (.044) |
| Posterior Thalamic Radiation (L) | PFBS | **0.007 [0.001, 0.012]** | .011 (.022) | -0.005 [-0.008, -0.001] | .014 (.032) | 0.001 [0.000, 0.001] | .022 (.075) | 6.075 | .048 (.081) |
| Superior Longitudinal Fasciculus (L) | PFBS | **0.005 [0.001, 0.008]** | .012 (.022) | -0.001 [-0.002, -0.000] | .036 (.060) | - | - | 4.396 | .036 (.069) |
| Superior Thalamic Radiation (L) | PFBS | **0.006 [0.002, 0.011]** | .003 (.009) | -0.004 [-0.008, -0.001] | .004 (.025) | 0.001 [0.000, 0.001] | .012 (.075) | **9.274** | .010 (.033) |
| Superior Thalamic Radiation (R) | PFBS | **0.006 [0.002, 0.010]** | .005 (.012) | -0.004 [-0.007, -0.001] | .019 (.040) | <0.001 [0.000, 0.001] | .044 (.088) | 6.267 | .044 (.078) |
| Uncinate Fasciculus (L) | PFOA | -0.003 [-0.006, -0.000] | .043 (.115) | 0.001 [0.000, 0.002] | .003 (.018) | - | - | **8.957** | .003 (.015) |
| Uncinate Fasciculus (L) | PFBS | **0.004 [0.001, 0.008]** | .026 (.041) | -0.001 [-0.002, -0.000] | .004 (.025) | - | - | **8.255** | .004 (.027) |
| Uncinate Fasciculus (R) | PFOA | -0.004 [-0.008, -0.000] | .030 (.115) | 0.004 [0.001, 0.007] | .009 (.033) | <0.001 [-0.001, -0.000] | .044 (.140) | **10.084** | .006 (.025) |
| Middle Cerebellar Peduncle | PFOA | **-0.009 [-0.014, -0.003]** | .002 (.012) | 0.005 [0.001, 0.009] | .028 (.069) | -0.001 [-0.001, 0.000] | .072 (.159) | 6.024 | .049 (.121) |
| Middle Cerebellar Peduncle | PFBS | **0.011 [0.004, 0.018]** | .001 (.007) | -0.006 [-0.011, -0.001] | .014 (.032) | 0.001 [-0.000, 0.002] | .051 (.092) | **8.273** | .016 (.039) |
| Forceps Major | PFOA | **-0.009 [-0.013, -0.005]** | <.001 (<.001) | 0.006 [0.003, 0.009] | <.001 (.004) | -0.001 [-0.001, -0.000] | .009 (.086) | 22.041 | <.001 (<.001) |
| Forceps Major | PFBS | **0.009 [0.004, 0.014]** | .001 (.005) | -0.005 [-0.008, -0.001] | .011 (.029) | 0.001 [0.000, 0.001] | .041 (.088) | **8.737** | .013 (.038) |
| Forceps Minor | PFOA | **-0.008 [-0.012, -0.004]** | <.001 (.005) | 0.006 [0.002, 0.009] | .001 (.011) | -0.001 [-0.001, -0.000] | .014 (.097) | **15.815** | <.001 (.003) |
| Forceps Minor | PFBS | **0.009 [0.003, 0.014]** | .001 (.007) | -0.006 [-0.010, -0.002] | .006 (.026) | 0.001 [0.000, 0.001] | .037 (.088) | **11.465** | .003 (.027) |

*Note:* Significant FDR-corrected associations of PFAS with fractional anisotropy (FA) in white matter tracts from longitudinal mixed models are reported above. The “PFAS B [95% CI]” term reflects the estimated effect and 95% confidence interval of a doubling in the given PFAS in the third column on FA at 4.5 years of age (i.e., the initial assessment). Uncorrected p-values (p) and FDR-corrected p-values (q) are reported for each estimate. Each model additionally included either a linear “PFAS x Age” interaction term or both a linear “PFAS x Age” and quadratic “PFAS x Age^2^” interaction term depending on whether modeling fit was improved when allowing for a nonlinear age trend. Models that were better fit linearly report dashes in the nonlinear columns. The significance of PFAS associations with age trends was determined from the likelihood ratio test (LRT) comparing model fit with and without the interaction of PFAS with linear and (if applicable) quadratic age terms. Bolded text indicates significant PFAS associations at baseline (“PFAS B [95% CI]”) or significant PFAS associations with age trajectories (LRT χ²) after FDR corrections.

**Sensitivity Analysis (S3.1):** PFAS associations with FA in white matter tracts after removing images with >3 mm motion

To preserve the largest sample possible, diffusion images were not excluded due to motion in the main analysis. Sensitivity analyses were conducted on a subset of participants with < 3mm motion to assess whether PFAS associations with FA in white matter tracts were independent of motion. Across all timepoints n=129 scans had > 3mm of motion (n_Y4.5_= 69, n_Y6_=47, n_Y7.5_=3, n_Y10.5_=10), resulting in a subset of 1011 observations from n=440 participants, compared to the larger sample of n=1140 observations from 455 participants. Analysis of PFAS associations with FA at baseline (age 4.5) and with longitudinal trajectories of FA across white matter tracts were repeated using the subset. As shown in Table S3.1 below, PFBS continued to be associated with higher FA in 13 of the 17 white matter tracts. However, PFOA remained associated only with lower FA in the middle cerebellar peduncle (below). In addition, PFNA, a long-chain PFAS, was associated with lower FA at 4.5 years of age in the left-hemisphere corticospinal tract. Together, these findings suggest that associations of PFBS with white matter microstructure at 4.5 years of age are largely independent of motion. Associations of PFOA with white matter microstructure largely do not survive FDR corrections when analyzing the subset with motion scores < 3mm. However, given the reduction in the sample size when evaluating the subset, it is possible that the attenuation of findings for PFOA is a consequence of reduced power to detect associations with PFOA exposure.

**Supplemental Table 3.1.** PFAS associations with FA in white matter tracts after removing images with >3 mm motion: significant associations at 4.5 years postnatal and links with age-trends

|  |  | **PFAS** |  | **PFAS x Age** |  | **PFAS x Age²** |  | **Trajectory** |  |
| --- | --- | --- | --- | --- | --- | --- | --- | --- | --- |
| **Tract** | **PFAS** | **B [95% CI]** | **p (q)** | **B [95% CI]** | **p (q)** | **B [95% CI]** | **p (q)** | **LRT χ²** | **p (q)** |
| Acoustic Radiation (R) | PFBS | **0.010 [0.005, 0.014]** | **<.001 (.002)** | -0.004 [-0.008, -0.001] | .009 (.071) | <0.001 [-0.000, 0.001] | .052 (.234) | **11.507** | .003 (.025) |
| Anterior Thalamic Radiation (L) | PFBS | **0.005 [0.001, 0.009]** | **.010 (.026)** | -0.002 [-0.005, 0.000] | .092 (.176) | <0.001 [-0.000, 0.001] | .276 (.473) | 6.811 | .033 (.139) |
| Cingulum (Cingulate Gyrus) (L) | PFBS | **0.008 [0.002, 0.014]** | **.008 (.023)** | -0.005 [-0.009, -0.001] | .010 (.071) | 0.001 [-0.000, 0.001] | .068 (.234) | **12.181** | .002 (.025) |
| Corticospinal Tract (L) | PFNA | **-0.009 [-0.014, -0.004]** | **.001 (.015)** | 0.006 [0.002, 0.010] | .001 (.037) | -0.001 [-0.001, -0.000] | .004 (.095) | 10.663 | .005 (.111) |
| Corticospinal Tract (L) | PFBS | **0.010 [0.005, 0.016]** | **<.001 (.002)** | -0.007 [-0.010, -0.003] | .001 (.018) | 0.001 [0.000, 0.001] | .003 (.056) | **11.815** | .003 (.025) |
| Corticospinal Tract (R) | PFBS | **0.009 [0.004, 0.015]** | **.001 (.008)** | -0.004 [-0.008, -0.000] | .029 (.123) | 0.001 [-0.000, 0.001] | .073 (.234) | 6.191 | .045 (.139) |
| Inferior Fronto-Occipital Fasciculus (L) | PFBS | **0.010 [0.004, 0.015]** | **.001 (.008)** | -0.004 [-0.008, -0.000] | .038 (.123) | <0.001 [-0.000, 0.001] | .169 (.312) | **9.774** | .008 (.041) |
| Inferior Fronto-Occipital Fasciculus (R) | PFBS | **0.005 [0.001, 0.009]** | **.023 (.047)** | -0.001 [-0.002, 0.000] | .109 (.196) | - | - | 2.575 | .109 (.172) |
| Inferior Longitudinal Fasciculus (L) | PFBS | **0.006 [0.001, 0.010]** | **.009 (.023)** | -0.003 [-0.006, -0.000] | .025 (.123) | <0.001 [-0.000, 0.001] | .057 (.234) | 6.003 | .050 (.139) |
| Posterior Thalamic Radiation (L) | PFBS | **0.006 [0.001, 0.011]** | **.017 (.038)** | -0.004 [-0.007, -0.000] | .036 (.123) | 0.001 [-0.000, 0.001] | .051 (.234) | 4.478 | .107 (.172) |
| Superior Longitudinal Fasciculus (L) | PFBS | **0.005 [0.001, 0.009]** | **.007 (.023)** | -0.003 [-0.005, -0.000] | .045 (.123) | <0.001 [-0.000, 0.001] | .132 (.312) | 6.512 | .039 (.139) |
| Superior Thalamic Radiation (L) | PFBS | **0.006 [0.003, 0.010]** | **<.001 (.003)** | -0.004 [-0.006, -0.001] | .001 (.018) | <0.001 [0.000, 0.001] | .005 (.056) | **11.182** | .004 (.025) |
| Superior Thalamic Radiation (R) | PFBS | **0.005 [0.001, 0.008]** | **.008 (.023)** | -0.002 [-0.005, 0.000] | .053 (.129) | <0.001 [-0.000, 0.001] | .088 (.234) | 4.105 | .128 (.193) |
| Middle Cerebellar Peduncle | PFOA | **-0.008 [-0.013, -0.004]** | **.001 (.018)** | 0.004 [0.001, 0.008] | .015 (.100) | -0.001 [-0.001, -0.000] | .038 (.186) | 6.873 | .032 (.170) |
| Forceps Major | PFBS | **0.006 [0.002, 0.011]** | **.003 (.014)** | -0.003 [-0.006, 0.000] | .063 (.141) | <0.001 [-0.000, 0.001] | .165 (.312) | 5.731 | .057 (.140) |

*Note:* Significant FDR-corrected associations of PFAS with fractional anisotropy (FA) in white matter tracts from longitudinal mixed models are reported above. The “PFAS B [95% CI]” term reflects the estimated effect and 95% confidence interval of a doubling in the given PFAS in the third column on FA at 4.5 years of age (i.e., the initial assessment). Uncorrected p-values (p) and FDR-corrected p-values (q) are reported for each estimate. Each model additionally included either a linear “PFAS x Age” interaction term or both a linear “PFAS x Age” and quadratic “PFAS x Age^2^” interaction term depending on whether modeling fit was improved when allowing for a nonlinear age trend. Models that were better fit linearly report dashes in the nonlinear columns. The significance of PFAS associations with age trends was determined from the likelihood ratio test (LRT) comparing model fit with and without the interaction of PFAS with linear and (if applicable) quadratic age terms. Bolded text indicates significant PFAS associations at baseline (“PFAS B [95% CI]”) or significant PFAS associations with age trajectories (LRT χ²) after FDR corrections.
